# ExocubeBio: an *in-situ* fluidic platform for microbial exposure on the International Space Station

**DOI:** 10.64898/2026.03.25.714121

**Authors:** David J. Burr, Ruben Nitsche, Elisa Ravaro, Severin Wipf, Pier Luigi Ganga, Michele Balsamo, Stefano S. Pellari, Francesco Caltavituro, Michael Gisi, Rodrigo Coutinho de Almeida, Pierfilippo Manieri, Antonella Sgambati, Claudio Moratto, Dennis J. Nürnberg, Adrienne Kish, Andreas Elsaesser

## Abstract

Space-based platforms currently represent the most accurate means to experimentally assess the influence of the space environment on biological systems. However, performing such experiments remains technically challenging and requires highly specialized instrumentation.

This study describes the current development and hardware qualification of ExocubeBio, a miniaturized experimental platform for *in-situ* biological space exposure. This experiment is scheduled for installation on the exterior of the International Space Station in 2027, as part of Exobio, the European Space Agency’s new generation exobiology exposure facility. ExocubeBio will expose live microbial samples to the low Earth orbit environment, and combine autonomous *in-situ* optical density and fluorescence measurements, with the capacity to return preserved samples to Earth. Achieving these experimental goals requires a specialized, robust and reliable hardware system.

The ExocubeBio hardware testing described here includes assessment of material biocompatibility and durability, functional validation of the miniaturized fluidic system, and optimization of the integrated optical subsystem for optical density and fluorescence measurements. These results demonstrate that the ExocubeBio experimental hardware components can each execute their core functional and operational requirements; subsystems allow for sample exposure, *in-situ* measurements of microbial cultures, and the chemical preservation of samples for post-flight analysis.

As ExocubeBio transitions from hardware development to mission readiness, the results presented here validate the overall design and engineering approaches utilized. By combining the strengths of *in-situ* monitoring and sample return, ExocubeBio represents a significant advancement in space-based experimentation, and will provide new insights into microbial responses to the space environment.

## 1. Introduction

Understanding how biological systems respond to the extreme conditions of space is a central focus of astrobiology and space life sciences. The unique environmental factors of space, including unfiltered solar ultraviolet (UV) radiation, galactic cosmic rays, vacuum, temperature cycling and microgravity [1, 2], are particularly challenging to experimentally replicate. While laboratory-based simulations on Earth form an indispensable foundation, they cannot fully reproduce the complex environmental interactions or the specific radiation environment beyond Earth’s atmosphere. In particular, the spectral shape and particle energy distribution of both solar UV and cosmic radiation differ significantly from those generated by terrestrial lamps, particle accelerators, or gamma sources [3, 4]. Likewise, synergistic interactions between radiation, vacuum, desiccation, temperature and microgravity are extremely difficult to mimic realistically on Earth [2, 5]. As such, while direct experimentation in space is logistically challenging, this is essential for obtaining biologically relevant data under accurate space-environment conditions.

Previous orbital and decent capsule exposure missions, such as Biopan [6], the EXPOSE-E [7, 8], -R [9] and -R2 [10] platforms, or Tanpopo [11] have revealed remarkable insights into the influence of true space conditions on the survival of microorganisms, the photochemical stability of biomolecules, and potential biosignature degradation pathways [3, 12-15]. However, such experiments function through passive sample exposure and rely solely on pre- and post-flight measurements. As such, transient, dynamic or rapid processes that occur during environmental exposure remain inaccessible. In contrast, free-flying nanosatellite missions such as GeneSat-1 [16], PharmaSat [17], SporeSat [18], O/OREOS [3] and recently BioSentinel [19], demonstrated the power of autonomous *in-situ* experimentation, including real-time measurements of active microbial growth and the photochemistry of organic molecules [4, 20-22]. However, these systems lack the capability for sample return, thus limiting their analyses to the miniaturized devices contained onboard.

The new European Space Agency (ESA)-funded experimental exposure platform, the Exobiology facility (Exobio; Figure 1a), is the first of its kind to combine miniaturized *in-situ* measurement technologies with sample return capabilities, thereby enabling comprehensive post-flight characterization. The goals of this platform address the strategic, technological and scientific priorities for astrobiology research in- and beyond low Earth orbit (LEO) outlined by both ESA [23, 24] and the National Aeronautics and Space Administration (NASA) [25]. The Exobio platform builds on both laboratory-based feasibility and hardware validation activities [26], and a dedicated technology demonstration operated aboard the International Space Station (ISS) for 132 days (SPECTRODemo,2019 [27]). This established the technical readiness and reliability of miniaturised spectroscopic instrumentation, fibre-optic switching, and detector performance for long-duration operation in LEO, thus providing critical heritage for the Exobio mission.

**Fig. 1:**
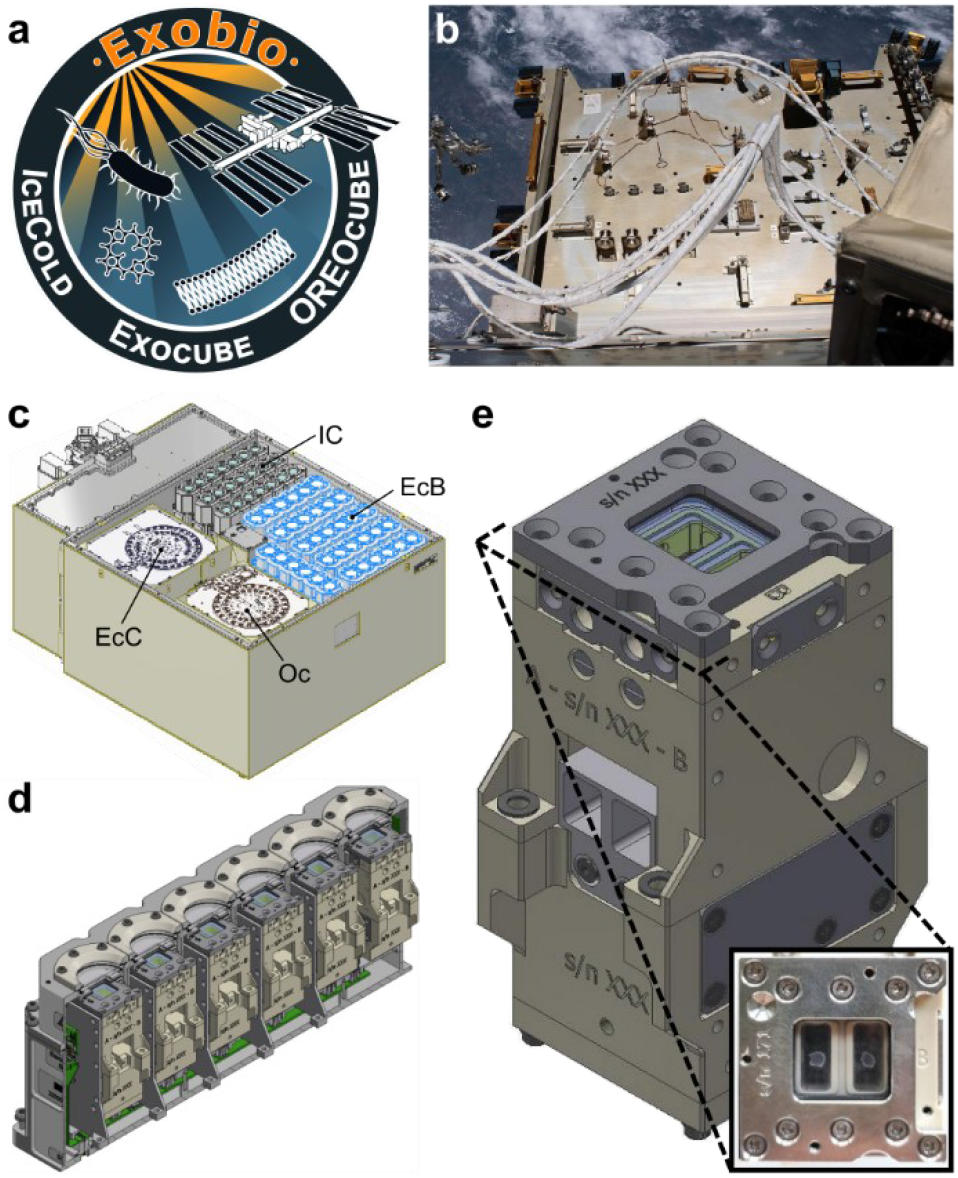
ExocubeBio design overview. **(a)** The Exobio project logo. **(b)** Photograph of the Bartolomeo science platform, outside of the ISS Columbus module (image credit: ESA/NASA, 2021 [28]). CAD illustrations depicting **(c)** the Exobio science module with hardware of each experiment labeled as follows: EcB = ExocubeBio (highlighted in blue), EcC = ExocubeChem, IC = IceCold, Oc = OREOcube, **(d)** a single ExocubeBio cartridge, shown in cross section to reveal the six experimental units contained inside, and **e**. a single ExocubeBio experimental unit. The inset is a top-down photo of an experimental unit, showing two independently housed, dry microbial samples deposited beneath the MgF_2_ window.

Exobio is currently in the final stages of development and is scheduled for installation on the exterior of the ISS in 2027. The facility will be mounted on the Bartolomeo platform (Figure 1b) outside of the Columbus module for 6-12 months. Bartolomeo provides power and data-uplink/downlink capabilities, and will orient the experiments in the forward-facing direction of the ISS, thereby minimizing the risk of accumulated contamination from outgassing, as observed on previous ISS exposure platforms mounted on the Zvezda module at the rear of the station [9]. During operation in the LEO environment, Exobio is expected to accumulate a total solar dose of ∼4920 MJ m^-2^. The platform integrates customized commercial-off-the-shelf (COTS) electronics and miniaturized spectrometer modules, whose manufacturing, benchmarking and environmental qualification (including vibration, thermal cycling and radiation tolerance) have previously been described [26].

Exobio comprises three independent but thematically complementary experiments: OREOcube, IceCold and Exocube, the latter of which consists of two distinct yet scientifically connected payloads, ExocubeBio and ExocubeChem (Figure 1c).

- ExocubeBio is a miniaturized *in-situ* biological exposure platform designed to host living microorganisms and monitor their response to space exposure conditions. It integrates controlled fluid handling with optical density (OD) and fluorescence measurements, to assess microbial growth and physiological responses under LEO conditions, with a major focus on the radiation environment.
- ExocubeChem will investigate the stability and transformation of organic molecules and biological membranes under prolonged LEO exposure. Using Fourier transform infrared (FTIR) spectrometry, this payload continuously monitors chemical modification induced by solar UV radiation and particle flux.
- IceCold concentrates on the combined biological influence of cold temperature and space radiation, in the context of the icy moons Europa and Enceladus. The growth of psychrophilic microorganisms will be actively monitored (via OD) during exposure to the LEO temperature oscillations and radiation environment.
- OREOcube will conduct an astrochemical investigation of biomolecules and organic-inorganic bilayers. UV-Vis absorption spectroscopy will be used to quantify radiation-induced chemical modifications of organic thin films and hybrid materials during space exposure.

Together these experiments constitute major advances in both space exposure technology and in furthering our understanding of how organisms and biomolecules endure, change with, or succumb to space conditions. In order to achieve these specific objectives however, each experiment must meet several technical requirements. The key objectives of ExocubeBio specifically are to observe microbial survival, recovery and damage induced by exposure to space radiation and growth in microgravity. To achieve these goals, ExocubeBio combines exposure of dried microbial samples in orbit, sample rehydration and *in situ* monitoring of microbial growth and metabolism, and finally the preservation of samples for the return to Earth. As such, there are a large and variable number of hardware requirements. This paper presents the validation and optimization of the experimental hardware that will be used to autonomously perform the ExocubeBio experiment.

ExocubeBio occupies a total volume of 45 L and is comprised of 11 sub-units termed cassettes (Figure 1d). Six cassettes will be exposed to solar radiation, while the remaining five will remain shielded for the duration of the experiment, functioning as in-flight dark controls. Each ExocubeBio cassette houses six experimental units (Figure 1e), each of which is equipped with dual fluidics and optical detection systems, thermal control, and the capacity to maintain an internal atmospheric pressure (whilst the exterior of the cassette is exposed to the vacuum of space). Each experimental unit houses two independent, dried microbial samples (Figure 1e, inset). In addition, a complete and matching hardware configuration will be utilized to carry out a ground reference control experiment, performed in parallel to the in-flight installation of ExocubeBio.

ExocubeBio will incorporate microorganisms relevant to astrobiology, aiming to address questions in radiation biology, planetary protection, life-support system development, planetary exploration and life-detection. Preliminary ground-based testing under experimentally simulated space stressors, including desiccation and UV irradiation, has identified several extremotolerant species suitable for flight integration. Possible candidate species span over a broad range of biological complexities, including radiation-resistant bacteria, halophilic archaea, and photosynthetic cyanobacteria or unicellular eukaryotic green algae. Model species from each of these clades provide the means to compare different survival strategies, DNA repair mechanisms, and oxidative stress mitigation. Additionally, ExocubeBio will also integrate protocellular samples, enabling the fundamental study of simple organized cellular components, such as lipid-encased genetic material. These function as an intermediary point of comparison between microbial astrobiology and organic astrochemistry studies [29].

By integrating real-time monitoring while in orbit, in combination with sample return across a diverse gradient of biological complexity, ExocubeBio represents a transformative next-generation astrobiology platform [24]. The simultaneous exposure of organisms to unfiltered space radiation and microgravity, followed by the *in-situ* monitoring of their physiological state will provide information unobtainable from passive exposure missions. The preservation and return of samples further expand the scientific potential of ExocubeBio, allowing for detailed post-flight analysis including microscopy, high-resolution spectroscopy, or potentially -omics analysis. The combination of *in-situ* data and post-flight analysis can provide unprecedented insights into physiological stress adaptations, subtle biomolecular changes, and the limits of life in space. However, such results are reliant on a robust and stringent hardware system.

Moving toward flight readiness, this paper presents the scientific qualification of the ExocubeBio hardware. These include assessing the biological compatibility and durability of the hardware components, functional validation of the autonomous miniaturized fluidic system, and the optimization of the optical detection system. This ensures a stable performance across the environmental conditions expected on the exterior of the ISS.

## 2. Methods

### 2.1 Experimental Design

This work describes the testing and validation of individual aspects of the ExocubeBio experimental hardware, each required to achieve the mission objectives. Each specific hardware test performed is related to a function of the ExocubeBio experiment, which following installation, will be conducted over three phases (Figure 2) during the operational period. The operational processes of each experimental phase highlight the essential function of the ExocubeBio hardware, and hence are the focuses of the validation testing described in this paper.

**Fig. 2:**
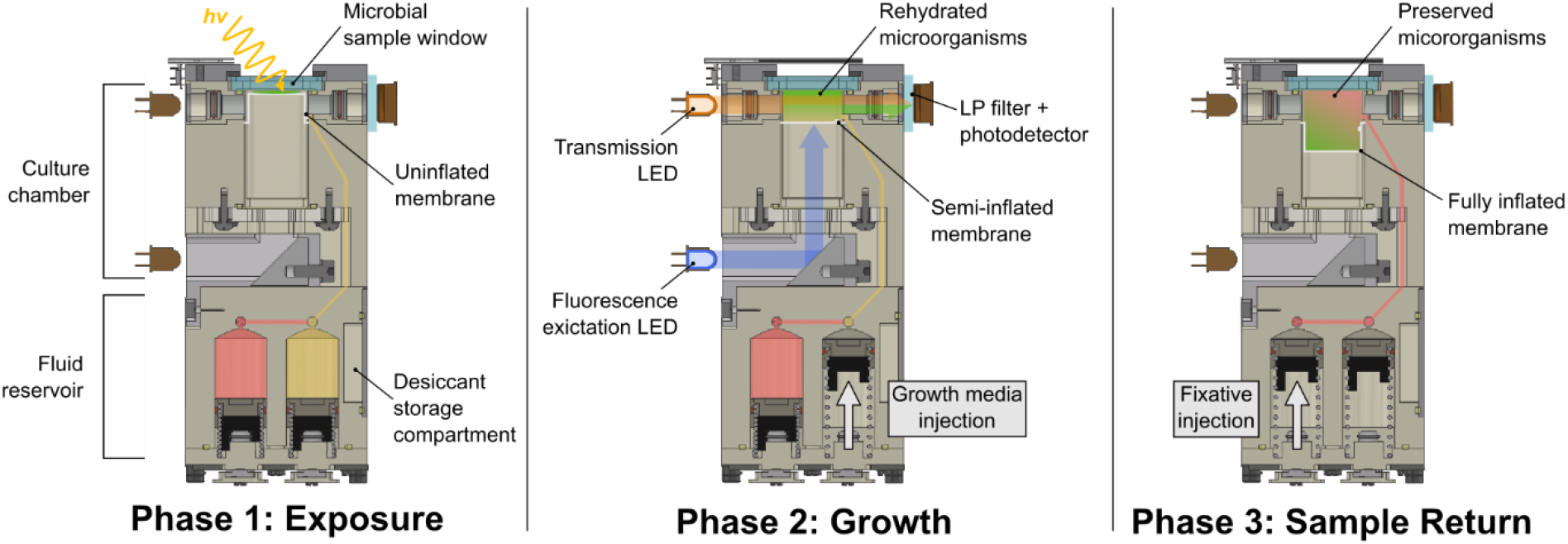
CAD diagrams illustrating the intended hardware function at each ExocubeBio experimental phase. As these diagrams are conceptual examples, some elements have been visually simplified or omitted for legibility. Illustrative elements or those not visible in this diagram include the liquid transport channels that run perpendicular to the plane of view shown, thermal plates housed on either side of each experimental unit, and air lines that connect the desiccant storage compartment to the gaseous headspace beneath the silicone membrane.

- Phase 1: Exposure A total of 128 live microbial samples across a range of biological complexities will be contained in experimental units installed outside of the ISS. Of these, 72 samples will be directly exposed to solar radiation for a range of durations, while the remaining 56 samples will remain continuously shielded, serving as dark controls. All microbial samples will be dried (either by aspiration or lyophilization) and housed between the MgF_2_ sample window and silicone membrane of each culture chamber (Figure 2, left). The highly UV-transmissive properties of MgF_2_ [30] will allow the solar-UV exposed samples to accumulate a species-dependent dose of solar radiation. It is of critical importance that samples are maintained in a dry state during exposure, as moisture not only influences microbial states of metabolic stasis [31], but also plays a key role in irradiation dynamics. For instance, the presence of water during UV exposure results in the formation of bio-degradative hydrogen peroxide, hydroperoxyl and hydroxyl radicals [32-34]. The time required to accumulate the specific solar-radiation dose is dictated by the orientation of the ISS in relation to the sun and will be measured via *in-situ* dosimetry acquired using an onboard UV-Vis spectrometer (Flame-T, Ocean Optics, Germany). Estimated solar dose rates have been modeled by simulations carried out by the Bartolomeo platform developer, Airbus Defence and Space (see Figure S1 and associated supplementary materials). The dark control samples will be consistently isolated from solar radiation, and are comprised of 30 in-flight, unirradiated microbial control cultures and 26 blank (uninoculated) media control samples. Each experimental unit is fitted with an Au-coated shutter above the MgF_2_ sample window. The shutters can be independently remotely triggered, enabling the solar exposure period to be terminated at different time points. This allows sample pairings within each experimental unit to accumulate independent solar radiation doses.
- Phase 2: Growth After concluding the exposure-phase, each culture chamber will be filled with a sub-milliliter volume of species-specific microbial growth media, injected from the first of two fluid reservoirs via the remote triggering of a spring-loaded piston. This fluid transfer is designed to operate reliably in microgravity, with the increase of volume in the culture chamber being accounted for by the expansion of the flexible silicone membrane (Figure 2, center). The gas-permeable nature of the silicone membrane allows for gas exchange with the headspace below. This, in combination with the temperatures control capabilities of each culture chamber (20 or 30°C ±3°C) allows growth of the surviving and rehydrated microorganisms. Each culture chamber is equipped with a miniaturized optical-detection system, comprised of a photodetector and either a 600 or 700 nm LED, with the latter also serving as a low-intensity growth light for photosynthetic samples (Figure 2; further detailed in Section 2.4). This allows periodic absorption measurements to be performed over time, providing a proxy measurement for the real-time, *in-situ* microbial growth of each sample. In addition, a secondary LED provides excitation light to the sample at 90° relative to the photodetector (reflected off a static 45° mirror; Figure 2), with wavelengths selected according to the spectral properties of specific fluorescent probes or intrinsic microbial fluorophores. Emitted fluorescence is recorded by the same photodetector. Therefore, ExocubeBio can simultaneously acquire *in-situ* information on cellular growth and metabolic processes.
- Phase 3: Sample Return Upon completion of the exposure phase, microbial activity can be remotely terminated through the injection of a chemical fixative. Fixation is intended to occur once cultures reach early-stationary phase, a time point selected to ensure sufficient biomass accumulation while minimizing variability associated with exponential growth or decline-phases stress responses. The addition of fixatives will halt cellular metabolism and preserve morphology for subsequent post-flight analyses. Fixative delivery is achieved via activation of each sample’s second fluid reservoir, which will inject a defined volume of fixative solution into the culture chamber. Here, the silicone membrane will expand to its engineered maximum displacement, allowing an approximate 1:1 mixing ratio within the culture chamber and maintaining sample containment (Figure 2, right). Following fixation, samples will be maintained at 2-8°C during deinstallation and return to Earth. Based on the operational requirements of each experimental phase, the following specific hardware functions were identified for testing: long-duration material functionality and biocompatibility testing; verification of the miniaturized fluidic system to provide sample isolation, rehydration and fixation; and quantified optimization of the absorption and fluorescence measurement system. It should be noted that less specific functions relating to the Exobio platform as a whole are not detailed in this article. These include vacuum containment, thermal isolation, and shock and vibration testing. Additionally, many of the COTS electronics have previously been environmental verified for space-flight use [26].

### 2.2 Component Validation

#### Biological compatibility testing

As maintaining active microbial cultures is a primary function of the ExocubeBio hardware, it is essential to ensure that microbial growth is not inhibited by any hardware components in contact with growth media. As such, biological compatibility testing (BCT) was conducted on each prospective hardware component that could be in contact with liquid growth media (during pre-flight and exposure-phases, as well as all components in the culture chamber). To do so, 16 unique components (Table S1) were sterilized (by autoclave or immersion in 70% ethanol), dried, and then independently submerged in one of six aqueous growth media solutions (Table 1). The volume of growth media was defined by the surface area of each hardware component, providing a similar surface area-to-volume ratio as the in-flight configuration (Table S1). All BCT solutions were stored in the dark, at 4°C, for 6 months. Additional samples of each growth medium (containing no hardware components) were prepared and stored in the same manner, acting as controls.

**Table 1:** Growth media and inoculation conditions used in BCT.

| Media | Ingredients or manufacturer | Inoculation species | Incubation conditions |
| --- | --- | --- | --- |
| GN101 | 4.28 M NaCl, 81mM MgSO<sub>4</sub>, 27 mM KCl, 10 mM 3Na-Citrate, 1% w/v Peptone Oxoid LP0034 (ThermoFisher Scientific, Germany)[35] | *Halobacterium salinarum* (ATCC 700922) | 37°C, 110 rpm orbital shaking |
| Tryptic Soy Broth (TSB) | 30 g L<sup>-1</sup> (Becton Dickinson, Germany) | *Bacillus subtilis* (DSM 10) | 30°C, 110 rpm orbital shaking |
| M9 minimal | 10.5 g L<sup>-1</sup> M9 Broth, 1.5 mM MgSO<sub>4</sub>, 0.1 mM CaCl<sub>2</sub>, 4.0 g L<sup>-1</sup> D-glucose (Merek, Germany) | *Escherichia coli* (DSM 498) | 37°C, 110 rpm orbital shaking |
| Luria/Miller LB Broth (LB) | 25 g L<sup>-1</sup> (Carl ROTH, Germany) | *Escherichia coli* (DSM 498) | 37°C, 110 rpm orbital shaking |
| BG-11 | 1x ready to use formulation (Life Technologies, Germany) | *Synechocystis* sp. (PCC 6803) | 28°C, 110 rpm orbital shaking, 50 µmol photons m<sup>-2</sup> s<sup>-1</sup> |
| TAP | 1x ready to use formulation (Life Technologies, Germany) | *Chlamydomonas reinhardtii* (CC-125 mt+) | 28°C, 110 rpm orbital shaking, 50 µmol photons m<sup>-2</sup> s<sup>-1</sup> |

Following the storage period, 4 mL subsamples of each growth media were aliquoted to 10 mL glass culture vials, and were inoculated with 100 µL of stationary-phase culture from the corresponding microbial model species (Table 1). To ensure the ExocubeBio hardware is compatible with a broad range of microorganisms, the five model species included representatives of archaea, Gram-positive bacteria, Gram-negative bacteria, cyanobacteria and photosynthetic microeukaryotes. Cultures were aerobically incubated (as detailed in Table 1) for a maximum duration of 14 days. Cellular growth was estimated via correlation with periodic OD measurements at either 600 or 730 nm (NanoPhotometer Classic, Implen, Germany).

#### Component appositeness

To ensure that exposure to the chemical compounds used in the experiment did not compromise the function of any hardware, 23 prospective components (Table S2) were independently submerged in four representative aqueous solutions. These included TSB and M9 minimal growth media (Table 1), each with 1.5% (v/v) DMSO (a common chemical used for dissolving potential fluorophores), an aldehyde-based fixative solution (25 mM HEPES buffer (adjusted to pH 7.5) + 0.5% (v/v) glutaraldehyde + 0.4% (v/v) paraformaldehyde), and phosphate-buffered saline (PBS) that served as a non-nutritive aqueous control matrix for material compatibility assessment. Triplicate components were independently stored under an incubation profile mimicking preparation, flight and installation times; 210 days at 8°C (±2°C), 60 days at 33°C (±0.5°C), 60 days at 8°C (±2°C). Following storage, the integrity of each component was validated through comparison against unexposed hardware, using three separate means: (i) measuring a change in component mass, (ii) visual inspection of material degradation or defects at x30 magnification (SmartScope MVP 250, OGP, Switzerland), and (iii) a leak-tightness test. The latter entailed fully assembling an experimental unit, integrating individual exposed components. The units were then filled and maintained under vacuum (pressure <10 mbar) for 60 min at 20°C. Leak tests were deemed successful when vacuum exposure resulted in <0.032 g mass loss.

Beyond general material assessment, the silicone membrane responsible for maintaining liquids within the culture chamber was specifically evaluated for its long-term functional integrity. This membrane serves two essential purposes within the ExocubeBio hardware; its mechanical flexibility allows the accommodation of varying liquid volumes during fluid injection, and the gas-permeable of the silicone membrane [36] enables passive O_2_ and CO_2_ exchange between the culture chamber and headspace. This gas diffusion supports aerobic metabolism and, in the case of photosynthetic organisms, provides access to CO_2_ required for active growth. However, as the silicone membrane is positioned directly beneath the MgF_2_ sample window (Figure 2), it will be exposed to the full solar spectrum during the exposure phase. To assess durability under these conditions, membranes were subjected to prolonged irradiation using a collimated xenon arc lamp solar simulator (UXL-302-0, Ushio, Germany; SciSun 300, Sciencetech, Canada), which approximates the spectral shape and intensity of the solar UV flux beyond Earth’s atmosphere [37, 38](Figure S2; 200-400 nm integrated intensity ≈ 100 W m^-2^). Following irradiation, membrane functionality was evaluated by integrating each UV-exposed silicone membrane into an experimental unit and inflating in with 1400 µL of water. Membranes were then monitored for 60 min for leakage or pressure loss.

Additionally, to validate the capacity of the silicone membrane to allow aerobic microbial growth, microorganisms were cultured in customized 20 mL gas-free glass vials, sealed with silicone membranes. The same model species were cultured as per the BCT (see Table 1), with *E. coli* only being cultured in LB media. Growth rates were compared against control aerobic cultures, incubated without sealing vials with a silicone membrane. As per BCT testing, periodic OD measurements at either 600 or 730 nm (NanoPhotometer Classic, Implen, Germany) were used to compare the growth rates of each sample.

### 2.3 Fluidics Functionality

#### Sample isolation and rehydration

To validate the efficacy of the silicone membrane in maintaining isolation between the dried microbial samples and the liquid contained in the liquid transport channels and reservoirs, the relative humidity was monitored over time. To remove any residual moisture, prior to assembly, all hardware components were dried under vacuum, at 60°C, for a minimum of 16 h. Each experimental unit was assembled with ∼450 mg of dry silica-beads (0.2-1 mm diameter, Merck, Germany) included within the desiccant storage compartment (Figure 2). The relative humidity was monitored using a custom digital humidity-monitor array in place of the sample window. This consisted of dual micro-humidity sensors (HIH120-021-001, Honeywell, Germany), independently epoxy-sealed in a custom 3D printed housing (BiomedClear Resin, Formlabs, USA) with the same dimensions as the MgF_2_ sample window. The lower face of each humidity sensor recess was left open, allowing the sensors to effectively reside in the same position within the culture chamber as the dried microbial samples in the flight configuration.

Humidity levels were compared between three different experimental unit configurations. A nominal flight-configuration unit was assembled with the liquid transport channels and reservoirs filled with a total of 2.39 mL of PBS. A wet treatment was created by perforating the push-fit plug of the silicone membrane with a 600 µm diameter hypodermic needle, resulting in a sub-mm hole between liquid transport channels and the culture chamber. Assembly and liquid filling were performed in the same manner as the nominal configuration. Finally, a dry experimental unit was assembled with a standard (un-perforated) silicone membrane, however the liquid transport channels and reservoirs were left unfilled, and the fluid ports and gas-exchange ports were left open to the environment. The gaseous headspace of all units was purged with dry air. Each unit was stored under a nitrogen atmosphere within a glovebox (MB-200B, M. Braun, Germany), maintained <0.1 ppm of atmospheric H_2_O, at room temperature, for over 1400 h (>60 days).

Following dry storage, samples must be autonomously rehydrated. To assess this capability, experimental units were prepared and filled in a nominal flight configuration, with microbial samples (∼10^7^ *E. coli* cells) air-dried onto the sample window within each culture chamber. The spring-loaded pistons of the fluid reservoirs were triggered, and rehydration was visually evaluated both top-down (through the sample window) and horizontally via the optical pathway. Inspections were made for both complete dissolution of the microbial samples and for the presence of trapped bubbles.

#### Sample fixation

To enable post-flight analysis of cellular phenotypic features, a defined volume of chemical fixative will be injected into each culture chamber at the conclusion of the growth phase. Fixatives must function (halting metabolism and preserving cellular ultrastructures) despite being diluted to levels that comply with health and safety standards for crewed spaceflight to the ISS, and being stored in the flight hardware for an extended period prior to use. As such, the functionality of multiple chemical-fixative solutions was compared. Four concentrations of aldehyde-based fixative (diluted in 100 mM cacodylate buffer, 88 mM sucrose, 1 mM CaCl_2_.H_2_O, pH adjusted to 7.4) were prepared: a standardized control concentration (2% paraformaldehyde (PFA), 2.5% glutaraldehyde (GA)), 0.5% total aldehydes (0.2% PFA, 0.25% GA), 0.1% total aldehydes (0.02% PFA, 0.025% GA), no aldehydes (all percentage values are v/v). These were compared against a formalin-free alternative, NOTOXhisto (NTH; Scientific Device Laboratory, USA), in undiluted and 50% (in PBS) concentrations.

The fixation capacity of the standard aldehyde concentration fixatives was also examined following exposure to a simulated 1-year dose of cosmic radiation levels onboard the ISS. 1.85 mL aliquots of fixative solution were stored in amber polypropylene microvials (ThermoFisher Scientific, Germany), and exposed to either a single 1.25 MeV step of Co-60 gamma radiation or 175 MeV of proton radiation. Radiation exposures were performed at the Fraunhofer Institute, Germany, and the Paul Scherrer Institute, Switzerland, respectively.

To assess the ability of different fixative treatments to preserve cellular ultrastructures, stationary phase *C. reinhardtii* (CC-125 mt+) was examined after fixation. *C. reinhardtii* was utilized as a model species due to its well documented subcellular morphology (that has been imaged through both transmission electron microscopy (TEM) [39], and with spectroscopic imaging techniques [40]), and its historic use in spaceflight experiments [41-44]. To do so, ∼10^9^ cells were resuspended in each fixative treatment solution and incubated at room temperature, in the dark for a minimum of 2 h. Post-fixation staining was performed with 1% (w/v) OsO_4_, and cells were dehydrated and embedded in Low Viscosity Resin (Agar Scientific, UK). Ultrathin sections (<100 nm) were cut from each sample, using an ultramicrotome (Reichert ultracut S, Leica, Germany) fitted with a diamond knife (Histo 45°, DiATOME, USA), and mounted on nickel square mesh grids (AGG2200N, Agar Scientific, UK). Subsequent cell staining was performed with UranyLess and 3% Reynolds lead citrate (Delta Microscopes, France). Cellular sections were imaged using TEM, with an electron energy of 120 kV, 60.80 µA beam current and a medium spot size (JEM-1400Flash, JEOL, Japan). Image stitching and brightness correction was performed with the TEM manufacturer software (TEM Center 1.7.18.2349, 2006-20018 JEOL, Japan).

### 2.4 Optical Quantification

#### Optical density measurements

The optical transmission detection system of the flight hardware consists of either a 600 nm (LED600 L, Thorlabs, Germany) or 700 nm LED (LED700-33au, Roithner LaserTechnik, Austria), supplying light that passes through the culture chamber (13.2 mm sample length). Transmitted light is then measured by a miniature photodetector (S2386-44K Si photodiode, Hamamatsu, Japan). The optical transmission system was validated via correlation, using varying concentrations of OD-standards (S0110710 Ink, Waterman, France) previously measured via UV-Vis spectrometry (7315 spectrophotometer, Jenway, Germany). Blank measurements were performed using experimental units filled with distilled water.

The raw voltage output of the photodetectors was first thermally corrected, using LED board-mounted temperature sensors. A common correction factor (1) was applied across matching LEDs.

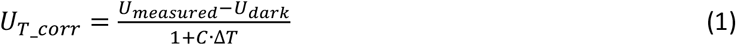

where Δ*T* is the difference between the temperature during measurement and the time zero temperature. *U*_*dark*_ is the voltage measurement made while the transmission LED is switched off. LED wavelength specific correction factors were *C*_*600 nm*_ = -0.0132 K and *C*_*700 nm*_ = -0.0229 K.

Values were converted to OD using the Beer-Lambert law (2), which also allowed for normalization against a typical 10 mm pathlength.

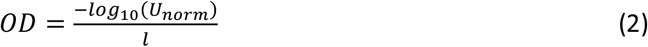

where *U*_*norm*_ is the *U*_*T_corr*_ value during measurement divided by the time zero *U*_*T_corr*_ value, and *l* is the pathlength.

#### Promotion of photosynthesis

Experimental units fitted with 700 nm LEDs are designed to house photosynthetic microorganisms. In addition to allowing transmission measurements, these LEDs must supply light sufficient for photosynthesis to occur. To validate this, comparative cultures of two photosynthetic model species, the cyanobacterium *Synechocystis* sp. (PCC 6803), and the eukaryotic microalga *C. reinhardtii* (CC-125 mt+) were grown under either red light supplied by the 700 nm LED (integrated intensity from 600-680 nm = 19.2 (±2.0) µmol photons m^-2^ s^-1^), or control white-light LEDs (W25E, ThorLabs, Germany; integrated intensity from 600-680 nm = 16.2 (±0.7) µmol photons m^-2^ s^-1^). With the exception of these modified light conditions, organisms were aerobically cultured under the conditions specified in Table 1. Growth was monitored via OD measurements at 730 nm (NanoPhotometer Classic, Implen, Germany).

#### Fluorescence optimization

The excitation light of the fluorescence-detection system is supplied at one of three selected wavelengths: 450, 507 or 522 nm (LED450-06; B5-433-B505, Roithner LaserTechnik, Austria; LTL2V3TGX3KS-032A, LITEON Technology, Taiwan). This is reflected off a custom-made 45° aluminum mirror, passes through the lower face of the translucent silicone membrane, and enters the culture chamber perpendicular to the photodetector. A wavelength-specific lowpass filter is housed between the culture chamber and the photodetector, blocking wavelengths below either 480, 520 or 540 nm. This minimizes the detection of scattered excitation light, whilst higher wavelength emitted light (and transmitted light) pass through the filter, reaching the photodetector. Excitation LEDs and lowpass filter combinations (including tested combinations that were not selected for implementation) are details in Table S3.

Fluorescent standard solutions, with an optimal excitation of 420 or 550 nm (F004 or F005, BAM, Germany), were utilized to validate the ExocubeBio fluorescence-detection system, both in the presence and absence of lowpass filters. Fluorescence spectra were acquired by exchanging the photodetector for a cosine-corrected optical fiber, attached to a calibrated UV-Vis spectrometer (HR4000, Ocean Insight, Germany), with a wavelength resolution of 0.25 nm. Using the ExocubeBio photodetectors, measurements were acquired once per second over a 30 s interval and corrected using the mean dark background measurement. The total detected light intensity was compared against scattered light intensities, by exchanging fluorescent standard solutions with 100% ethanol. The emitted fluorescent signal was defined as the difference between the total detected light intensity and the scattered light intensity. A signal-to-noise ratio was derived, defining the noise as the standard deviation of the fluorescent signal.

### 2.5 Statistical Analysis

BCT results were assessed by determining the difference in the mean cell number of each treatment from that of each respective positive control, at four microbial growth-phases (early exponential, late exponential, early stationary and late stationary). Cell numbers were enumerated via correlation with OD measurements (at either 600 or 730 nm for non-photosynthetic and photosynthetic organisms, respectively), with mean cell counts determined through use of biological replicates. The mean difference in cell number across all four timepoints was compared using a single-tailed, one sample t-test at the 95% confidence level. BCT induced changes in cell number were also compared against positive controls at each specific time point using a one-way analysis of variance (ANOVA).

Similarly, a one-way ANOVA was used to compare total detected fluorescence intensities against scattered light intensities. A regression analysis was used to compare OD measurements made with the experimental hardware against reference spectrometer measurements. The variation of the fit of correction used is expressed as a coefficient of determination (R^2^).

All statistical analyses were performed with RStudio (v. 2023. 12.0+369, Posit Software, USA). Spectra peak integration was performed using OriginPro (v. 2023, OriginLab Corporation, USA).

## 3. Results

### 3.1 Hardware Validation

#### Biological compatibility testing

Of the 16 prospective hardware components examined, only two demonstrated a consistent inhibitive effect on microbial growth across multiple species: ethylene-propylene diene monomer (EPDM) TA 50-75 and TA 50-60 (Table 2). More specifically, EPDM TA 50-75 treated media resulted in a >95% loss in viability for four of the five tested microbial species. EPDM TA 50-60 also was detrimental to the growth of multiple species, but has the largest effect on *H. salinarum*, causing a 98% loss in viability.

**Table 2:**
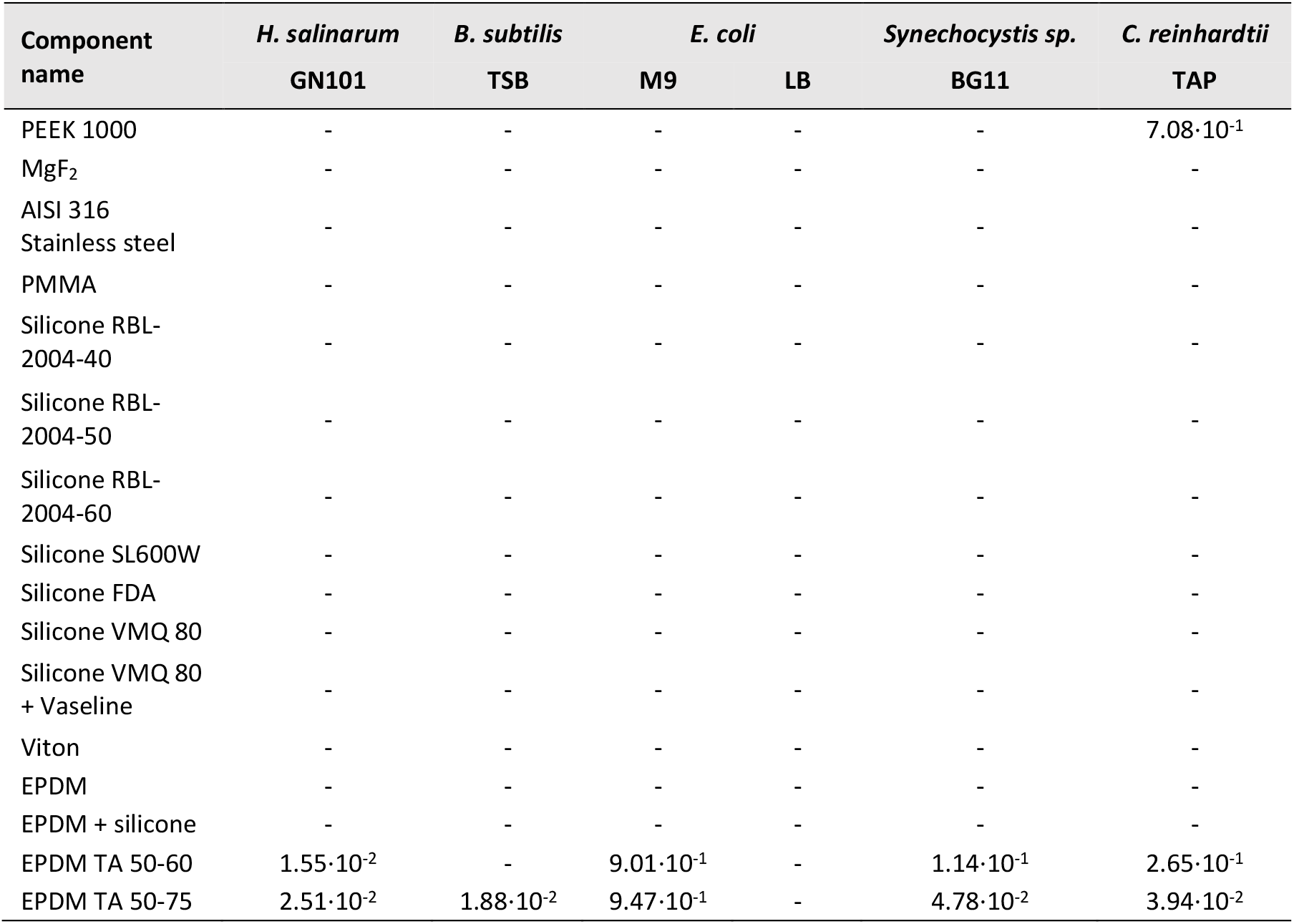
BCT results. Mean survival fraction (N/N_0_) of microbial cultures growing in media exposed to prospective hardware components (n = 3, biological replicates). Values are shown for treatments that resulted in a statistically significant decline at the 95% confidence level, whereas the “–” symbol represents no significant reduction in survival (as determined by a single-tailed, one sample t-test).

*E. coli* demonstrated the highest resilience to both EPDM TA 50-75 and TA 50-60, however the extent of the resilience was influenced by the specific growth media which the components were stored in. Although treatments were identically inoculated, the detrimental effects of EPDM TA 50-75 and TA 50-60 were more pronounced in M9 minimal media cultures than LB media treatments. This may be due to either the LB media having a protective property or damaging compounds more readily leached from the components into the M9 minimal media.

Aside from the EDPM-based treatments, exposure to other prospective hardware components did result in some growth fluctuations, though these reductions were generally sporadic. For instance, the maximum growth of *C. reinhardtii* cultures was periodically moderately reduced by several specific components (Table S1), but when averaged over time this difference was only apparent in PEEK 1000 exposed media (Table 2).

Stored media controls did not induce any reduction in growth, regardless of species. Based on the results of the BCT, EPDM TA 50-75 and TA 50-60 were excluded from use in ExocubeBio. All remaining components were approved for use.

#### Component appositeness

Aqueous-solution exposure had no major influence on the structure or function of all tested prospective hardware components, and no major differences occurred between different solutions. Across all tested solutions, the mean induced change in mass proportion of non-lubricated components was -0.58% (±0.008%). The mean mass of lubricated components reduced by 5.28% (±0.01%; Table S2), likely due to loss into solution. Despite the small amount of mass loss observed, no optical artefacts were introduced, and sealing interfaces performed robustly under vacuum exposure. Specifically, no significant material degradation or defects were observed under magnified inspection, with the only observed change being a small accumulation of oxidation on selected aluminum components. Additionally, all functionality tests performed with exposed hardware components all passed, with units remaining leak-tight following 60 min under 1 bar delta pressure.

In contrast, accumulated solar UV radiation had a significant impact on membrane performance. Silicone membranes retained structural integrity and functionality following exposure to cumulative UV doses up to 36 MJ m^-2^, with all tested samples passing the 60-minute leakage assessment described above. Exposure to 44 MJ m^-2^ resulted in the formation of micro-fissures in some membranes, with one of the eight samples failing the leakage test. At cumulative UV radiation doses of 55 MJ m^-2^ all tested membranes became brittle and exhibited mechanical failure during functionality testing. Based on these results, the UV dose accumulated during the exposure phase must remain below 36 MJ m^-2^ to ensure maintained sealing performance within the hardware configuration.

All aerobic species examined demonstrated growth when cultured in vessels sealed with silicone membranes. However, the presence of membranes reduced the maximum optical density reached by all species (Figure S3a-e). This species-specific reduction in accumulated biomass was likely the result of limited gas exchange associated with the membrane barrier. Despite this reduction, measurable growth occurred for each species. The silicone membranes were therefore suitable for implementation in the ExocubeBio experimental hardware.

### 3.2 Fluidics Functionality

#### Sample isolation and rehydration

The nominal ExocubeBio fluidics configuration maintained a relative humidity in the culture chamber between 10-15% over the entire >1400 h measurement period (Figure 3a, orange). Unfilled units left open to the dry environment of the glovebox demonstrated a consistent decline in relative humidity, decreasing from the ∼10% observed upon sealing the unit, towards a complete lack of atmospheric moisture (Figure 3a, yellow). These two treatments are contrasted by the experimental units prepared with a perforated silicone membrane. The sub-mm hole in the push-fit plug resulted in an immediate and rapid increase in relative humidity, exceeding 90% by 260 h (Figure 3a, blue).

**Fig. 3:**
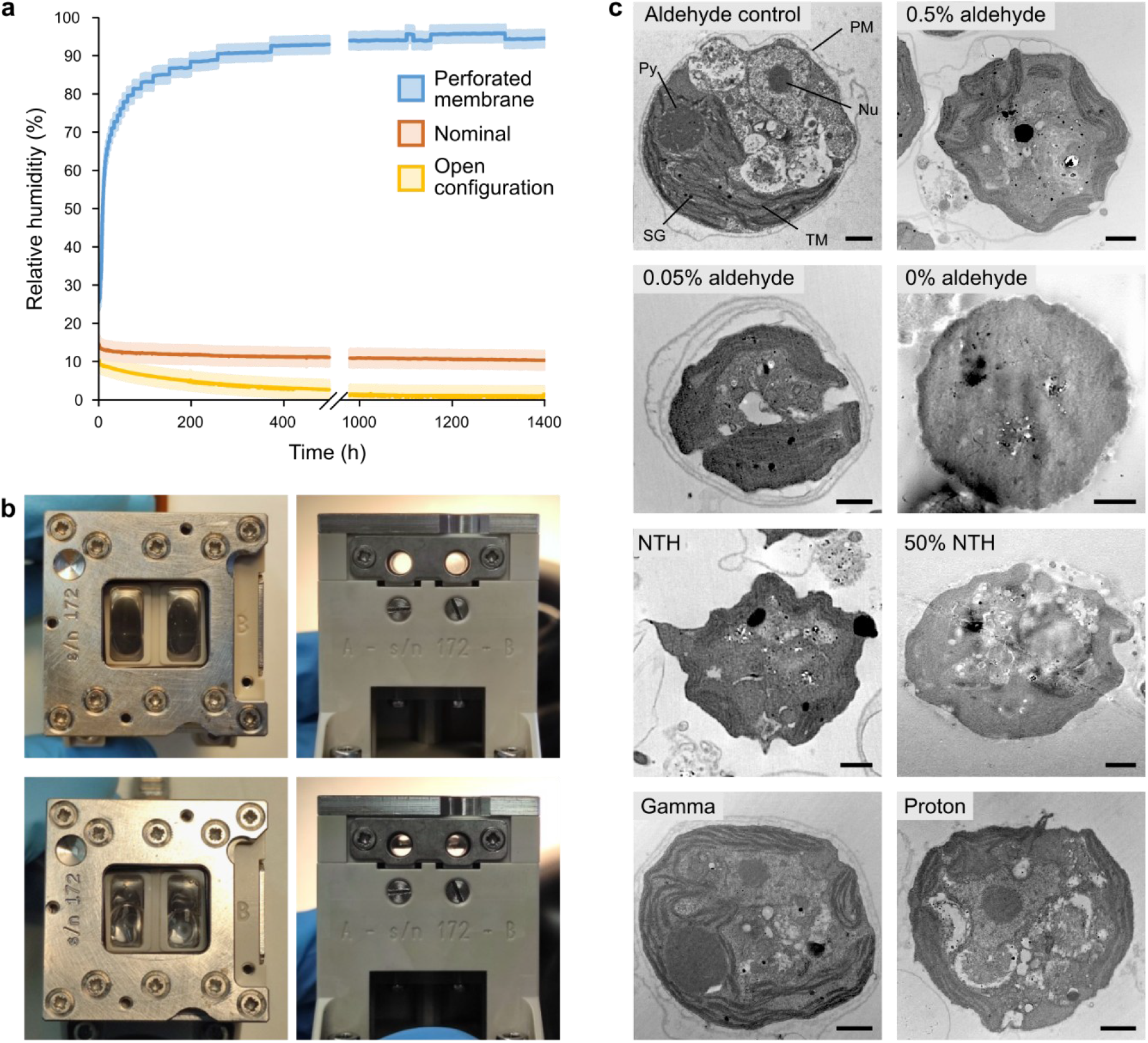
ExocubeBio fluidic system functionality. **(a)** Mean relative humidity measured over time, within Exocube culture chambers (n = 2, biological replicates). The nominal hardware configuration (orange) is compared against experimental units prepared with a perforation in the silicone membrane between the culture chamber and the liquid transport channels (blue), and a unit prepared without liquids and left open to the dry, nitrogen atmosphere (yellow). Shaded regions represent standard deviations plus humidity sensor accuracy specifications. **(b)** Photos of liquid-filled culture chambers of an ExocubeBio experimental unit, shown from top-down (left) and through the optical pathway (right). The upper pair of photos demonstrate a nominal activation of the fluidics system, completely filling the culture chamber. In contrast, the lower pair of photos are a representative example of a partial experimental failure, due to large bubbles within the culture chamber, indicative of incorrect filling. **(c)** TEM imagining of *C. reinhardtii* cells preserved with different fixative solutions, including four concentrations of aldehydes (4.5% control – 0%), undiluted and a 50% dilution of the aldehyde-free fixative NOTOXhisto (NTH), and standard aldehyde-based fixative solutions exposed to simulated ISS-level, annual doses of either gamma or proton irradiation. Labeled cellular features are as follows: Nu = nucleus (within the nucleolus), PM = plasma membrane, Py = pyrenoid, SG = starch granule, TM = thylakoid membrane. Scale bars equal 1 µm.

These results demonstrate that nominal assembly of the ExocubeBio experimental units allows the culture chambers to successfully remain sealed from the external environment. Additionally, the silicone membrane (in combination with the silica beads housed in the desiccant compartment) is sufficient to keep the microbial samples dry during transport, installation and the exposure phase. Following this point, growth media will be injected into the culture chamber, dislodging the push-fit plug of the silicone membrane, rehydrating surviving microorganisms and beginning the growth phase.

Under nominal conditions, triggering of the fluid reservoir pistons resulted in a rapid injection of liquid into the culture chamber. Dried microbial samples were immediately rehydrated, with visible dissolution completely occurring in <2 min (Figure 3b, top). However, repeated operation revealed that incorrect filling of the fluid reservoirs or liquid transport channels may result in bubbles trapped within the culture chamber (Figure 3b, bottom). As these have the potential to hinder sample rehydration or obscure the optical pathway, exclusion of air bubbles from the culture chamber is of critical importance. This requirement is made particularly stringent by conditions in microgravity where bubbles cannot escape through the gas-permeable silicone membrane by buoyancy. Unwanted air can be excluded during the filling process due to several specific design choices. These include minimizing the number of angles within the fluid reservoirs and liquid transport channels, and incorporating passive valve structures and gas exclusion outlets. As such, nominal filling and function should allow for fluid movement into the culture chamber with minimal to no initial bubble formation (Figure 3b, top).

#### Sample fixation

Both the chemical composition and fixative concentration were largely influential on cellular preservation, as expected. *C. reinhardtii* cells fixed with an aldehyde control solution (∼4.5% total aldehyde concentration) demonstrated good maintenance of cellular shape and the plasma membrane. Intracellular ultrastructures were also well preserved. Visible organelles include the nucleus within the nucleolus, tubules penetrating the pyrenoid, distinct stacks of thylakoid membranes and starch granules within the chloroplast, and other subcellular compartmentalization within the cell (Figure 3c, aldehyde control).

Decreasing the aldehyde concentration resulted in a stepwise reduction in preservation capacity. Fixing *C. reinhardtii* in a 0.5% total aldehyde solution decently maintained ultrastructure preservation, with major membrane-bound organelles still being visible. However, this did result in a moderate loss of cell shape, with the cell wall retracting from the plasma membrane (Figure 3c, 0.5% aldehyde). A further reduction in aldehyde concentration continued to emphasize this morphological degradation. The 0.05% aldehyde concentration resulted in minimal ultrastructure preservation, with the thylakoid being the only visible organelle. Near-complete separation of the plasma membrane also occurred. Without the presence of any aldehydes, no visible plasma membrane could be identified, and there was a complete loss of ultrastructure preservation within the cell (Figure 3c, 0.05% aldehyde and 0% aldehyde).

The formalin-free fixative, NTH, yielded poor cellular preservation both at a 50% dilution and when used at full strength. Undiluted NTH resulted in comparable *C. reinhardtii* ultrastructure preservation as observed with the 0.05% aldehyde solution, while 50% NTH was most similar to cells fixed without aldehydes. Both NTH solutions lead to a near-complete separation or total loss of the plasma membrane, and a major degradation in cell shape (Figure 3c, NTH and 50% NTH).

Doses of either gamma or proton radiation simulating 1-year of accumulated cosmic radiation onboard the ISS, had no apparent degrative effect on the preservative properties of aldehydes. Observed cellular features, ranging from the cell shape to the intracellular ultrastructures of *C. reinhardtii* cells fixed with irradiated aldehyde solutions are highly comparable to those of the aldehyde control (Figure 3c, gamma and proton). This indicates that an aldehyde-based fixative is more appropriate for use than NTH during the ExocubeBio experiment.

### 3.3 Optical Quantification

#### Optical density measurements

OD measurements made with the ExocubeBio optical system were correlated against a reference benchtop spectrophotometer. Both the 600 and 700 nm light sources required correction, with a second order polynomial yielding a particularly strong fit (both R^2^ > 0.997; Figure 4a). With this quadratic correction applied, the accuracy can be expressed as the mean difference in measured and reference OD values. For the 600 and 700 nm light sources, this delta was 0.015 and 0.013, respectively. Similarly, the highest standard deviation at any measured point can be used to represent the precision of the optical systems. For the 600 nm light source this value was 0.03, whereas the 700 nm light was 0.02.

**Fig. 4:**
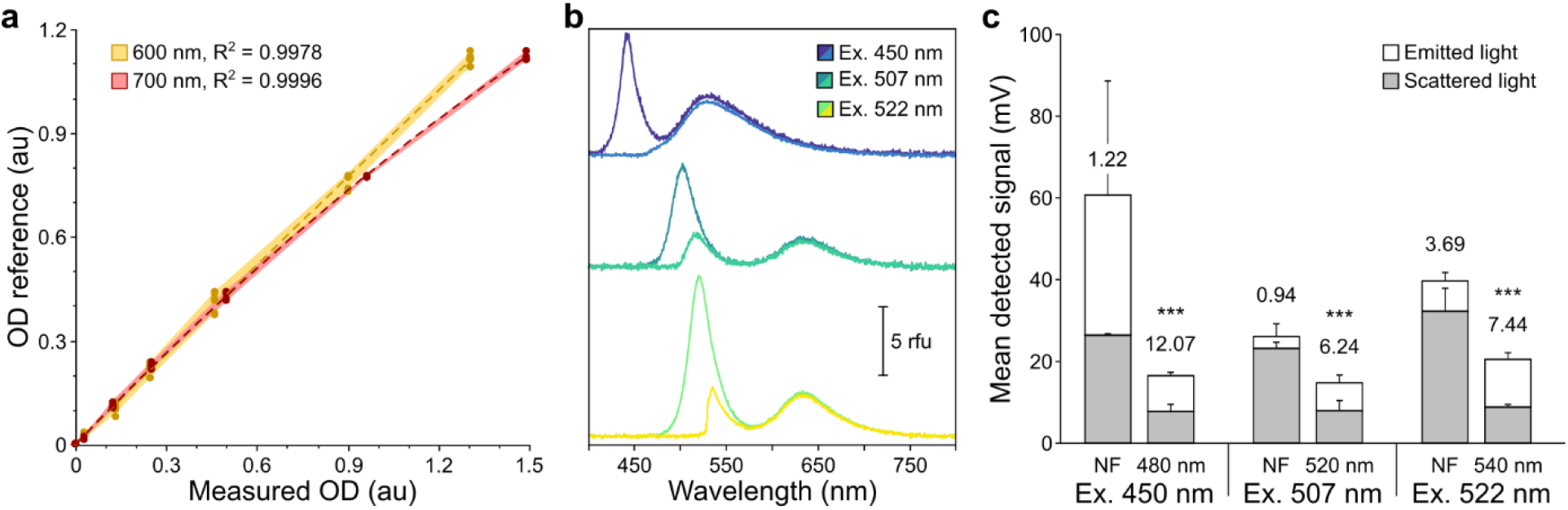
ExocubeBio optical detection system quantification. **(a)** Mean correlative measurements of OD signals acquired with either 600 or 700 nm light sources (depicted in orange and red, respectively), compared against signals of a reference spectrometer (n = 4, biological replicates). Shaded regions represent standard deviations. Dashed lines represent a quadratic fit, with the coefficient of determination (R^2^) denoted for each. **(b)** Fluorescence spectra (in relative fluorescence units) of fluorescence standards (BAM F004 or F005) measured under different ExocubeBio optical detection configurations. Excitation wavelengths include 450 (blue), 507 (green-blue) and 522 nm (lime green). Measurements are shown in the absences (dark shades) and the presence (light shades) of respective lowpass filters. **(c)** Mean light intensities measured under different ExocubeBio optical detection configurations (n = 6, biological replicates). The x-axis lists treatment pairs of supplied excitation wavelengths (450, 507 and 522 nm) and the absence (NF) or presence of a lowpass filter (480, 520, 540 nm). The total detected light of each treatment is divided into the emitted fluorescence signal (white) and scattered light (grey). The signal-to-noise value of each treatment is listed, and asterisks denote a statistically significant difference (p < 0.01) between the total mean light intensity and the mean scattered light. Error bars represent standard deviation.

#### Promotion of photosynthesis

The red light supplied by the 700 nm transmission LED was sufficient to allow growth of both cyanobacteria and unicellular algae. When compared against white-light cultures, growth under red light resulted in slower exponential growth of *Synechocystis* sp. (Figure S3d) and a slight reduction in the maximum growth obtained by *C. reinhardtii* (Figure S3e). However, as these induced changes were minimal and could be accounted for by comparison against control samples, the 700 nm light source was considered adequate for the cultivation of photosynthetic microorganisms in the ExocubeBio hardware.

#### Fluorescence optimization

The use of lowpass filters in the ExocubeBio fluorescence-detection system minimized scattered excitation light, while allowing the detection of the emitted fluorescence signal. When measured spectroscopically, the three selected lowpass filters (480, 520, 540 nm) resulted in a reduction of detected excitation intensities by 95.3, 66.8 and 78.9% (integrated between 415-480, 450-570 or 470-580 nm, respectively; Figure 4b). In contrast, the fluorescence signal was only minimally affected by the lowpass filters, with the mean change in detected emission intensity (integrated from either 480, 570 or 580 to 750 nm) being -9.2% (±0.8%; Figure 4b). Spectra of only the ethanol backgrounds further demonstrates the reduction in scattered light due to the use of lowpass filters (Figure S4).

The improvement brought about by the lowpass filters can also be seen through measurements made with the ExocubeBio photodetectors. Across all three selected excitation LEDs, there was a large reduction in the detected scattered light when lowpass filters were present. While this caused a reduction in the total detected light, proportionally the emitted fluorescence signal increased (Figure 4c). Additionally, a lowpass filter dramatically reduced the particularly high variation in the total detected signal from the 450 nm excitation LED (Figure 4c). As such, incorporation of the lowpass filters into the ExocubeBio optical setup resulted in a 9.9-, 6.7- and 2.0-fold improvement in the signal-to-noise ratio of the 450, 507 and 522 nm excitation LEDs, respectively.

Additional combinations of excitation LEDs and lowpass filters were also tested. 352 and 558 nm excitation LEDs were insufficiently bright to differentiate the total detected light from dark background measurements, and thus were excluded from further analysis. Several further combinations of excitation LED and lowpass filter also yielded promising fluorescence excitation spectra (Figure S4), and the scattered light measured with the ExocubeBio photodetector could often be statistical differentiated from total light intensity (Table S4). However, the specific combinations of excitation LED and lowpass filter selected for use in the ExocubeBio optical detection system offer the highest available signal-to-noise (Table S4), and wavelengths appropriate for the excitation of various biologically relevant fluorescent compounds.

## 4. Discussion

The primary objective of the ExocubeBio developmental testing was to validate that the specialized hardware meets all functional and operational requirements, in order to perform a fully autonomous, microbiological exposure and culture experiment on the exterior of the ISS. The results presented here demonstrate that the ExocubeBio hardware has the capacity to execute the operations required by each experimental phase: exposure, growth and sample return. Broadly, material components were confirmed as biologically compatible and functionally durable; fluidic subsystems were validated for reliable sample isolation, rehydration and preservation; and the optical detection system was optimized for sample interrogation via consecutive absorption and fluorescence emission detection.

BCT demonstrated that two potential gasket materials, EPDM TA 50-60 and TA 50-75, were inhibitive to a variety of microbial model species. As the foundational material (vulcanized rubber) did not result in microbial growth inhibition, the altered ethylene, monomer and oil ratios required to change the material Shore A hardness value (as indicated by the TA nomenclature) caused an undefined, broadly bio-interfering compound to leach from these components into aqueous growth media. This excluded these materials from incorporation into the ExocubeBio in-flight hardware configuration. It should be noted that the BCT results presented here only documented components that negatively influenced the growth of the tested model organisms, due to introduced toxicity. Some hardware components resulted in a slight increase in the microbial growth of specific species. For instance, the o-ring and gasket material, Viton, affected GN101 growth media causing an increase in the growth of *H. salinarum*. However, minor hardware-induced discrepancies from normal growth rates will not alter the scientific outcome of the experiment, as all in-flight samples will be compared directly against ground controls exposed to identical hardware components.

Prolonged exposure of hardware components to various experimentally relevant solutions (growth media, buffer or fixative) did not induce any consequential changes. However, extended doses of simulated solar UV-visible spectrum eventually resulted in the gas-permeable silicone membrane becoming brittle and unfunctional. Although the planned ∼6-month mission duration for the Exobio platform as a whole, will greatly exceed the maximum UV-irradiation time of the silicone membranes, the ExocubeBio Au-coated shutters will be closed after no more than 100 h, blocking further incoming solar radiation. Thus, the silicone membrane UV-life span shall not be exceeded. Taken together with the BCT results, this material-testing dataset serves as a valuable resource for the design of experimental hardware for future microbiology space-exposure missions.

Qualification of the fluidics system was performed over three steps: (i) maintaining sample dryness during exposure, (ii) rehydrating microorganisms at the beginning of the growth phase, and (iii) the delivery of chemical fixative into the culture chamber to preserve samples post-flight analysis. The silicone membrane (in combination with the stored silica desiccant) adequately isolated dry samples from the fluid-filled liquid transport channels, with only a ∼10% increase in relative humidity from complete dryness. Subsequently, samples were successfully rehydrated. However, this process was dependent on the exclusion of air bubbles during filling of the fluidics system. While the aerobic culture of microorganisms may result in microbubbles forming within the culture chamber (particularly during incubations performed in microgravity), we estimate this influence to be less consequential than larger trapped bubbles. As such, correct filling of the fluidics system represents a critical step in the success of the ExocubeBio experiment.

Due to the unconventional nature of this experiment, functional validation was performed at aldehyde concentrations below typical dilutions and following exposure to ionizing and particle radiation. Aldehyde-preservation capacity was directly correlated to concentration, but left unimpacted from radiation exposure. In contrast, the aldehyde-free fixative, NOTOXhisto, did not adequately preserve the tested microbial samples. As such, the ExocubeBio experiment will utilize an aldehyde-based fixative solution. Although PFA rapidly acidifies in the presence of atmospheric oxygen, degrading its preservation capacity [45], this issue is addressed by the design of the fixative reservoirs, which inherently exclude gas. In order to comply with astronaut health and safety requirements, aldehyde concentrations within the fixative reservoirs must be reduced below conventional levels. However, adequate microbial sample preservation was possible with a total aldehyde concentration as low as 0.5%. Previous studies have demonstrated that similar minimum aldehyde concentrations are required for the fixation of whole mammalian-cells, but this capacity is further improved by prolonged exposure times [46]. This suggests that the extended storage of ExocubeBio samples during the return phase may further aid preservation.

Despite the unavoidable design limitations due to the inherent volume constraints of ExocubeBio’s miniaturized optical system, the transmission LED-photodetector architecture yielded accurate and precise OD measurements with either 600 or 700 nm light sources. This indicates that the optical configuration is sufficient to monitor microbial growth up to OD = 1.4. Additionally, the 700 nm transmission LED fulfilled its secondary function, providing sufficient light for the photosynthetic growth of cyanobacteria and unicellular green algae.

While the integration of specific lowpass optical filters resulted in a significant improvement in the system’s ability to discriminate fluorescence emission from scattered background excitation light, this came at the cost of an overall reduction of detected intensity. This highlights the primary limitation of the ExocubeBio optical system; an inherent miniaturization-defined design constraint is the use of a single photodetector for both OD and fluorescence measurements. As these two measurement types result in vastly different light intensities reaching the photodetector, an extraordinarily wide dynamic range is required. As such, the fluorescence sensitivity of the ExocubeBio optical system did not reach the originally targeted performance levels. However, the fluorescence capability remains scientifically valuable, and the quantitative validation presented here will directly inform the selection of suitable fluorophores or fluorescent organisms for the flight experiment. Future fluorescence-based experimental platforms can further improve on these outputs through the use of separate and specifically tunned photodetectors for OD and fluorescence measurements.

This data confirms the core functionality of each ExocubeBio hardware subsystem required for planned *in-situ* experimentation in space. These outcomes validate the engineering decisions taken during development, and support the readiness of the instrument for integrated in-flight environmental testing. Now, as ExocubeBio enters its final preparation and integration phase, the project is transitioning from hardware development to mission readiness. In parallel to production of the final flight hardware, biological payload optimization is progressing. This includes the final selection and preparation of the microbial sample species, appropriate growth media and fluorescent probes. Together, these ongoing activities form the final set of steps required to prepare for flight integration, prior to launch.

The complimentary experiments that comprise the Exobio facility address many of the astrobiology and astrochemistry related goals and milestones outlined by ESA and NASA [23-25], and represents a significant advancement in experimental space exposure platforms. In particular, integrating both miniaturized *in-situ* monitoring and sample return allows for correlative analysis that will provide new insights into biomolecular stability, the limitations and origins of life, guide life-detection strategies, and provide essential knowledge for sustaining biological systems during long-duration human exploration.

## 5. Conclusion

The hardware subsystems that comprise ExocubeBio have demonstrated functional performance and technical readiness for autonomous *in-situ* microbiological space exposure experiments as planned for the Exobio mission. Material validation, fluidic system testing, and optimization of the optical detection subsystem indicate that the ExocubeBio experimental hardware can execute each critical operational step under conditions representative of LEO. Design elements such as the gas-permeable silicone membrane and integrated thermal control are intended to support aerobic microbial cultivation *in situ*. The *in-situ* monitoring capabilities of ExocubeBio, including optical density measurement and selective fluorescent detection, provide a framework for investigating time-dependent microbial responses to space exposure. These capabilities are complimented by the capacity to chemically preserve samples for return to Earth, enabling correlation of real-time measurements with advanced post-flight analyses. Together, this combined approach is expected to support investigations into space-induced physiological responses, metabolic transitions, dormancy processes and DNA repair dynamics. Beyond its application in LEO, the development of ExocubeBio establishes technological foundations for future biological space exposure platforms, including missions operating in more distant or deep-space environments.

## Supporting information

ExocubeBio Supplementary Information

## Acknowledgements

The authors gratefully acknowledge the European Space Agency for funding and supporting the development of the Exobio facility and for enabling its operation on the International Space Station. We further acknowledge Airbus Defence and Space for their contributions to the estimations of the solar dose received by Exobio, and for contributing critical information regarding the orientation and operational characteristics of the Bartolomeo platform. The authors also acknowledge Pascal Hass from the Department of Veterinary Medicine, Freie Universität Berlin, Germany, for performing the counterstaining and TEM imaging of thin-sectioned cellular samples.

## 6.2 Author Contributions

The scientific objectives of the ExocubeBio project were originally conceived by A.E., with A.K. and A.E. being responsible for related funding acquisition and scientific supervision. D.J.B. wrote and coordinated the original manuscript, with contributions from R.N., E.R., S.W., P.L.G., M.B., S.S.P, F.C., M.G., R.C.d.A., D.J.N, A.K. and A.E. Data collection and analysis were primarily performed by D.J.B., with additional contributions from R.N., E.R., S.W., M.B., S.S.P. and M.G. The design and manufacturing of the experimental hardware was performed by P.L.G., M.B. and S.S.P., with F.C. and M.G. being responsible for the optical system. R.C.d.A, A.S., P.M. and C.M. were responsible for project management, logistics and coordination. All authors contributed to the experimental design and approval of the final manuscript.

## 7. Data availability

The data used to support the findings of this study are available upon reasonable request. However, some material and component specifications are proprietary to Kayser Italia S.r.l. or OHB SE, and thus disclosure must be evaluated on a case-by-case basis.

