## Supplementary material for "ExocubeBio: an *in-situ* fluidic platform for microbial exposure on the International Space Station": ExocubeBio Supplementary Information

The expected annual solar dose accumulated by ExocubeBio was estimated through simulations carried out by Airbus Defence and Space as part of the Bartolomeo Exobiology Obscuration Analysis (document number: BTL-EXPO-ADSB-AN-000001, issue 2, 2023.12.04). Using an experimental unit model, a rectangular simulation sensor (typically used for satellite or aircraft field of view modeling) was orientated perpendicular to the sample-window boresight. The sensor field of view is constrained by four planes, defined by half angles in the YZ and XZ planes (Figure S1). The sample window sits at the vertex of these planes. Due to the highly UV-transmissive properties of $MgF_2$ [1], this has minimal to no influence on the accumulated solar dose reaching the microbial samples.

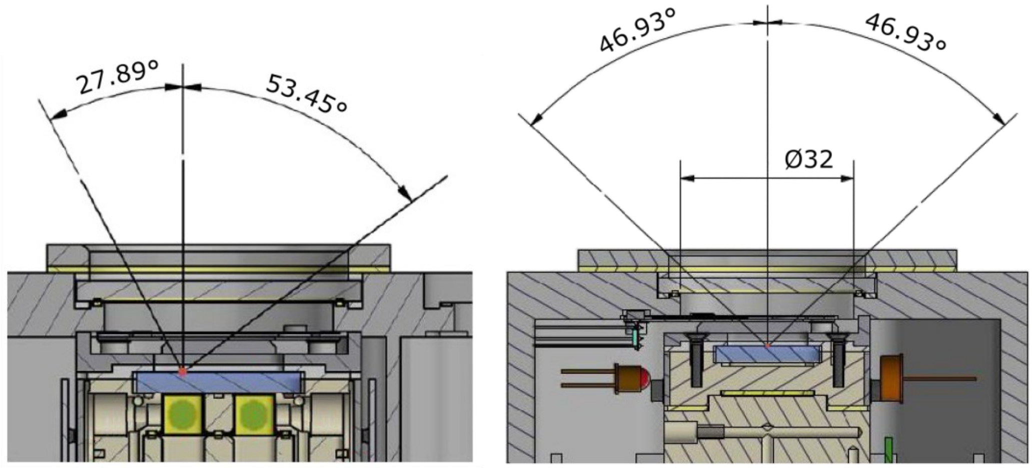


**Fig. S1: Sensor half angles utilized for ExocubeBio obscuration analysis.**
CAD diagrams are shown in the YZ section plane (left) and the XZ section plane (right). Illustration adapted from Bartolomeo Exobiology Obscuration Analysis (document number: BTL-EXPO-ADSB-AN-000001, issue 2, 2023.12.04), ©Airbus Defence and Space.

Lighting time analyses were conducted with a 60 s time step, using the most common ISS attitude, as-flown from mid-2018 to mid-2021 (180.1 days, yaw = -4.0°, pitch = -2.0°, roll = 0.7°). The orbit is based on the two-line element set from the ISS as initial value, and is then calculated using a simplified general perturbations propagator. This takes into account the effect of perturbations caused by the Earth's shape, drag, radiation and gravitational effects.

The solar flux ($H$) was calculated as follows (S1):

$$H = E_0 \cdot \sum(|\cos \alpha i| \cdot \Delta ti) \cdot QC_{avg} \tag{S1}$$

with a total solar irradiance ($E_0$) = 1367 W $m^{-2}$ [2], taking into account large solar panel rotation, $QC_{avg}$ as the average unobscured quotient of the field of view, $\alpha i$ denoting the sun incidence angle, $\Delta ti$ corresponding to simulation step size, and the sum including all time points ($i$), where the sensor has sun access.

The annual solar dose was calculated as (S2):

$$Solar_{total\ dose} = Solar\ flux \cdot t \cdot \left(1 - mean(obscuration\ \%)\right) \tag{S2}$$

where the average solar flux over the entire electromagnetic spectrum was 4.9212 MJ $m^{-2}$ $h^{-1}$ (multiplying $E_0$ by 3600, to convert from seconds to hours). The ExocubeBio zenith orientation varied from 3.6-5.1%, with a mean obscuration value of 3.8%. This resulted in an annual accumulated access duration of 1427.96 h. Therefore, the total incoming annual solar dose (over the entire electromagnetic spectrum) of ExocubeBio is estimated at 5656.1 MJ $m^{-2}$.

**Table S1: Prospective hardware components and BCT results over time.**

Component name abbreviations include polyetheretherketone (PEEK), American Iron and Steel Institute (AISI), polymethyl methacrylate (PMMA), Food and Drug Administration (FDA), ethylene-propylene diene monomer (EPDM). The supplied component surface area to liquid volume ratio is proportionally compared against the flight hardware ratio. Mean survival fraction ($N/N_0$) of microbial cultures growing in media exposed to hardware components (n = 3), at four time points. Values are shown for treatments that resulted in a statistically significant decline at the 95% confidence level, whereas the "–" symbol represents no significant reduction in survival (as determined by a one-way ANOVA). Light grey and dark grey shading represent a survival reduction of >1 log and >2 log, respectively.

| Component | Function | Δ surface area:volume ratio (%) | GN101 + *H. salinarum* | | | | TSB + *B. subtilis* | | | | M9 + *E. coli* | | | | LB + *E. coli* | | | | BG11 + *Synechocystis sp.* | | | | TAP + *C. reinhardtii* | | | |
|---|---|---|---|---|---|---|---|---|---|---|---|---|---|---|---|---|---|---|---|---|---|---|---|---|---|---|
| | | | Early exp. | Late exp. | Early stat. | Late stat. | Early exp. | Late exp. | Early stat. | Late stat. | Early exp. | Late exp. | Early stat. | Late stat. | Early exp. | Late exp. | Early stat. | Late stat. | Early exp. | Late exp. | Early stat. | Late stat. | Early exp. | Late exp. | Early stat. | Late stat. |
| PEEK 1000 | Experimental unit body | -0.0001 | - | - | - | - | - | - | - | - | - | - | - | - | - | - | - | - | - | - | - | - | - | - | $6.41 \cdot 10^{-1}$ | $4.56 \cdot 10^{-1}$ |
| $MgF_2$ | Sample window | 0.0016 | - | - | - | - | - | - | - | - | - | - | $8.45 \cdot 10^{-1}$ | - | - | - | - | - | - | - | - | - | - | - | $7.32 \cdot 10^{-1}$ | $5.15 \cdot 10^{-1}$ |
| AISI 316 Stainless steel | Valves | -0.0026 | - | - | - | - | - | - | - | - | - | - | - | - | - | - | - | - | - | - | - | - | - | - | - | - |
| PMMA | Bottom window | -0.0049 | - | - | - | - | - | - | - | - | - | - | - | - | - | - | - | - | - | - | - | - | - | - | - | - |
| Silicone RBL-2004-40 | Culture chamber membrane | -0.0005 | - | - | - | - | - | - | - | - | - | - | - | - | - | - | - | - | - | - | - | - | - | - | - | - |
| Silicone RBL-2004-50 | Culture chamber membrane | -0.0005 | - | - | - | - | - | - | - | - | - | - | - | - | - | - | - | - | - | $7.03 \cdot 10^{-1}$ | - | - | - | - | - | - |
| Silicone RBL-2004-60 | Culture chamber membrane | -0.0002 | - | - | - | - | - | - | - | - | - | - | - | - | - | - | - | - | - | - | - | - | - | - | - | - |
| Silicone SL600W | O-rings, gaskets | 0.0037 | - | - | - | - | - | - | - | - | - | - | - | - | - | - | - | - | - | $7.41 \cdot 10^{-1}$ | - | - | - | - | - | - |
| Silicone FDA | O-rings, gaskets | -0.0020 | - | - | - | - | - | - | - | - | - | - | - | - | - | - | - | - | - | - | - | - | - | - | $6.28 \cdot 10^{-1}$ | $5.05 \cdot 10^{-1}$ |
| Silicone VMQ 80 | O-rings, gaskets | 0.0000 | - | - | - | - | - | - | - | - | - | - | - | - | - | - | - | - | - | $7.41 \cdot 10^{-1}$ | - | - | - | - | - | $4.84 \cdot 10^{-1}$ |
| Silicone VMQ 80 + Vaseline | Lubricated O-rings, gaskets | 0.0000 | - | - | - | - | - | - | - | - | - | - | - | - | - | - | - | $9.38 \cdot 10^{-1}$ | - | - | - | - | - | - | $6.57 \cdot 10^{-1}$ | $4.98 \cdot 10^{-1}$ |
| Viton | O-rings, gaskets | 0.0000 | - | - | - | - | - | - | - | - | - | - | - | - | - | - | - | $9.33 \cdot 10^{-1}$ | - | - | - | - | - | - | $5.71 \cdot 10^{-1}$ | $5.63 \cdot 10^{-1}$ |
| EPDM | O-rings, gaskets | 0.0000 | - | - | - | - | - | - | - | - | - | - | - | - | - | - | - | - | - | - | - | - | - | - | - | - |
| EPDM + silicone | Lubricated O-rings, gaskets | 0.0000 | - | - | - | - | - | - | - | - | - | - | - | - | - | - | - | - | - | - | - | - | - | - | $7.82 \cdot 10^{-1}$ | $5.42 \cdot 10^{-1}$ |
| EPDM TA 50-60 | O-rings, gaskets | 0.0069 | $2.90 \cdot 10^{-2}$ | $1.58 \cdot 10^{-2}$ | $2.19 \cdot 10^{-3}$ | $1.49 \cdot 10^{-2}$ | $6.44 \cdot 10^{-1}$ | $8.29 \cdot 10^{-1}$ | $6.94 \cdot 10^{-1}$ | - | $8.23 \cdot 10^{-1}$ | $8.73 \cdot 10^{-1}$ | $7.90 \cdot 10^{-1}$ | - | - | - | - | - | $2.03 \cdot 10^{-1}$ | $7.50 \cdot 10^{-2}$ | $1.79 \cdot 10^{-1}$ | LETHAL | - | $2.43 \cdot 10^{-1}$ | $1.69 \cdot 10^{-1}$ | $1.83 \cdot 10^{-1}$ |
| EPDM TA 50-75 | O-rings, gaskets | 0.0014 | $4.35 \cdot 10^{-2}$ | $3.16 \cdot 10^{-2}$ | $2.08 \cdot 10^{-2}$ | $4.47 \cdot 10^{-3}$ | $3.94 \cdot 10^{-2}$ | $9.11 \cdot 10^{-3}$ | $1.37 \cdot 10^{-2}$ | $1.28 \cdot 10^{-2}$ | $8.03 \cdot 10^{-2}$ | $6.77 \cdot 10^{-1}$ | $7.38 \cdot 10^{-1}$ | $7.22 \cdot 10^{-1}$ | $4.98 \cdot 10^{-1}$ | $6.49 \cdot 10^{-1}$ | $6.89 \cdot 10^{-1}$ | $9.52 \cdot 10^{-1}$ | $1.10 \cdot 10^{-1}$ | $4.38 \cdot 10^{-2}$ | $1.29 \cdot 10^{-2}$ | $2.45 \cdot 10^{-2}$ | $5.60 \cdot 10^{-2}$ | $4.95 \cdot 10^{-3}$ | $3.45 \cdot 10^{-3}$ | $9.33 \cdot 10^{-2}$ |

**Table S2: Proportional change in prospective hardware component mass following prolonged aqueous solution exposure.**

| Component | Function | Δ mass (%) | | | |
|---|---|---|---|---|---|
| | | TSB media + 1.5% DSMO | M9 minimal media + 1.5% DSMO | Aldehyde fixative solution | PBS |
| PEEK 1000 | Experimental unit body | -0.04% | -0.09% | -0.07% | -0.05% |
| $MgF_2$ | Sample window | -0.05% | -0.04% | -0.04% | -0.04% |
| AISI 316 Stainless steel | Valves, screws and washers | -0.06% | -0.06% | -0.12% | -0.03% |
| PMMA | Bottom window | -3.13% | -3.11% | -3.24% | -3.25% |
| Silicone RBL-2004-40 | Culture chamber membrane | -0.24% | -0.39% | -0.21% | -0.18% |
| Silicone SL600W | O-rings, gaskets | -0.70% | -0.47% | -0.48% | -0.62% |
| Silicone FDA | O-rings, gaskets | -0.95% | -0.64% | -0.69% | -0.65% |
| Silicone FDA + Vaseline | Lubricated O-rings, gaskets | -4.67% | -5.49% | -2.99% | -4.05% |
| Silicone VMQ 80 | O-rings, gaskets | -1.11% | -0.70% | -0.88% | -0.70% |
| Silicone VMQ 80 + Vaseline | Lubricated O-rings, gaskets | -3.72% | -4.55% | -4.95% | -5.60% |
| Viton | O-rings, gaskets | -0.40% | -1.31% | -0.65% | -0.78% |
| Viton + Vaseline | Lubricated O-rings, gaskets | -6.32% | -3.56% | -3.24% | -4.76% |
| EPDM | O-rings, gaskets | -0.81% | -1.11% | -0.51% | -0.91% |
| EPDM + silicone | Lubricated O-rings, gaskets | -6.91% | -6.25% | -6.10% | -7.00% |
| EPDM TA 50-60 | O-rings, gaskets | -0.56% | -0.94% | -2.87% | -1.53% |
| EPDM TA 50-60 + silicone | Lubricated O-rings, gaskets | -7.17% | -5.96% | -6.87% | -5.36% |
| EPDM 70 Shore A UBA black | Gaskets | -0.50% | -0.35% | -0.31% | -0.35% |
| FKM - 75 SH A | Gaskets | -0.35% | -0.16% | -0.12% | -0.36% |
| Polycarbonate | Cartridge | -0.41% | -0.19% | -0.021% | -0.14% |
| Aluminum alloy 6061-T6 | Cartridge | -0.09% | -0.08% | -0.38% | -0.45% |
| Aluminum alloy 6061-T6 Lanthane conversion | Cartridge | -0.09% | -0.23% | -0.30% | -0.05% |
| Aluminum alloy 6082-T6 Lanthane conversion | Cartridge | -0.07% | -0.08% | -0.08% | -0.03% |
| Aluminum alloy 6082-T6 Sulfuric anodization, MILA-8625 Type I Class1 | Cartridge window | -0.04% | -0.23% | -0.03% | -0.90% |

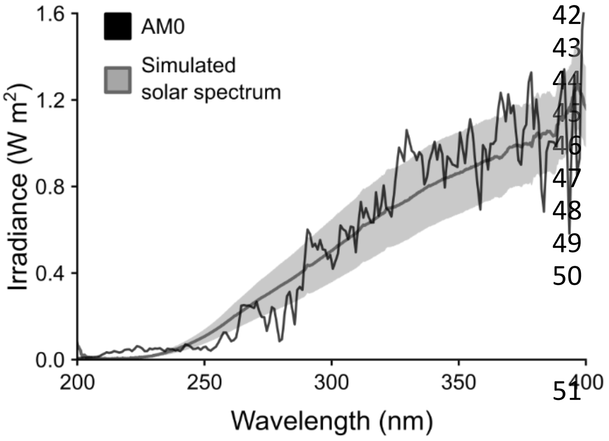


**Fig. S2: Mean simulated solar UV spectrum supplied, compared against AM0 UV spectrum.** The integrated intensity between 200-400 nm for the AM0 spectrum (black) = 106.59 W m$^{-2}$ [3, 4], and for the mean supplied UV spectrum = 102.68 ± 23.25 W m$^{-2}$. The mean supplied spectrum was determined from multiple measurements (n = 16, technical replicates) in the X-Y plane. The shaded region represents the standard deviation.

**Table S3: Details of fluorescence excitation and lowpass filters.**
Excitation LED and lowpass filter combinations selected for integration in ExocubeBio are italicized.

| Excitation wavelength (nm) | LED manufacturer | Lowpass filter wavelength (nm) | Lowpass filter manufacturer |
|---|---|---|---|
| 352 | FG350-R5.5WC015, The Fox Group, USA | 400 | FGL 400, Thorlabs, Germany |
| *450* | *LED450-06, Roithner LaserTechnik, Austria* | *480* | *Hoya Y48 #18-863, Edmund Optics, USA* |
| | | 515 | FGL 515, Thorlabs, USA |
| | | 525 | #15-212, Edmund Optics, USA |
| 490 | LED 490L, Thorlabs, Germany | 525 | #15-212, Edmund Optics, USA |
| *507* | *B5-433-B505, Roithner LaserTechnik, Austria* | 515 | FGL 515, Thorlabs, USA |
| | | *520* | *Hoya Y52 #18-871, Edmund Optics, USA* |
| 515 | WL-TMRW THT, Würth Elektronik, Germany | 520 | Hoya Y52 #18-871, Edmund Optics, USA |
| *522* | *LTL2V3TGX3KS-032A, LiteOn, Germany* | *540* | *Hoya O54 #66-040, Edmund Optics, USA* |
| | | 570 | FGL 570, Thorlabs, Germany |
| 558 | HLMP-K640, Broadcom, USA | 570 | FGL 570, Thorlabs, Germany |

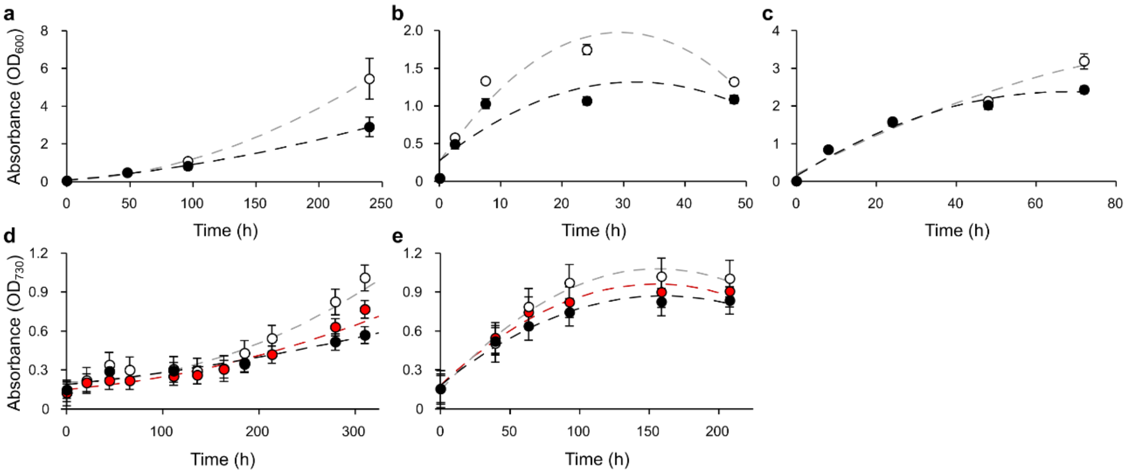


**Fig. S3: Microbial growth curves.**

Mean growth of **(a)** *H. salinarum*, **(b)** *E. coli*, **(c)** *B. subtills*, **(d)** *Synechocystis* sp. and **(e)** *C. reinhardtii* cultures (n = 4, biological replicates). Microorganisms were cultured under either aerobic control conditions (white), or while sealed with a silicone membrane (black). Photosynthetic microorganisms were either supplied with white light (during the two treatments specified above), or red light during an additional aerobic incubation (red). The extended lag-phases observed in *H. salinarum* and *Synechocystis* sp. are likely due to cultures being performed in small-volume culture vials, limiting gas exchange. Dashed lines show exponential fits, and error bars represent standard deviations.

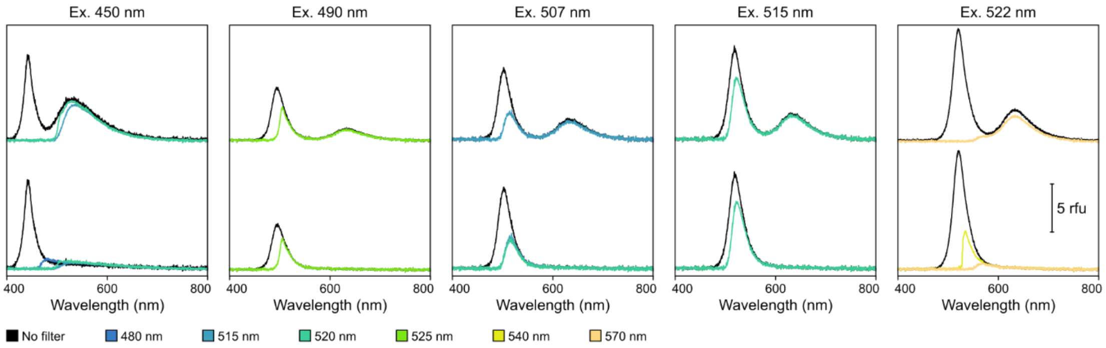


**Fig. S4: Fluorescence spectra of non-selected, optical detection configurations.**

Spectra (in relative fluorescence units) of BAM 004 or 005 fluorescence standards (top) and 100% ethanol backgrounds (bottom). Excitation wavelengths are titled, and lowpass filters are denoted in the legend.

**Table S4: Outputs and statistical analysis of fluorescence detection system configurations.** The mean total and scattered light intensities are listed for each excitation wavelength and lowpass filter combination, with their corresponding standard deviation in parentheses. Additionally, the derived signal-to-noise ratio (S:N) and results of each one-way ANOVA are listed. Excitation LED and lowpass filter combinations selected for integration in ExocubeBio are italicized.

| **Value** | **Ex. 450 nm** | | | | **Ex. 490 nm** | | **Ex. 507 nm** | | | **Ex. 515 nm** | | **Ex. 522 nm** | | |
|---|---|---|---|---|---|---|---|---|---|---|---|---|---|---|
| | **NF** | **480 nm** | **515 nm** | **525 nm** | **NF** | **525 nm** | **NF** | **515 nm** | **520 nm** | **NF** | **520 nm** | **NF** | **540 nm** | **570 nm** |
| Mean total light intensity (mV) | 60.78 (27.98) | 16.12 (1.63) | 14.57 (1.83) | 14.87 (1.79) | 16.94 (7.18) | 15.72 (4.42) | 27.05 (3.80) | 16.23 (3.57) | 13.45 (3.82) | 35.54 (4.61) | 27.12 (4.78) | 39.71 (4.44) | 20.52 (2.14) | 13.59 (2.61) |
| Mean scattered light intensity (mV) | 26.44 (0.32) | 7.76 (1.72) | 6.11 (1.31) | 5.82 (0.80) | 12.74 (3.47) | 9.23 (1.70) | 23.23 (1.49) | 10.82 (2.43) | 7.99 (2.43) | 26.31 (3.48) | 18.29 (3.55) | 32.30 (5.56) | 8.83 (0.61) | 4.55 (0.83) |
| S:N | 1.22 | 12.07 | 12.16 | 4.42 | 0.81 | 1.77 | 0.94 | 3.49 | 6.24 | 2.96 | 2.68 | 3.69 | 7.44 | 4.62 |
| $F_{1, 10}$ | 2.26 | 56.00 | 71.08 | 106.02 | 1.38 | 9.36 | 5.31 | 14.95 | 23.44 | 12.79 | 11.01 | 5.43 | 137.84 | 54.41 |
| p-value | 0.183 | $3.76 \cdot 10^{-5}$ | $7.42 \cdot 10^{-6}$ | $1.22 \cdot 10^{-6}$ | 0.267 | 0.012 | 0.044 | 0.003 | 0.001 | 0.005 | 0.008 | 0.042 | $3.59 \cdot 10^{-7}$ | $2.38 \cdot 10^{-5}$ |


[1] Heath DF, Sacher PA. Effects of a simulated high-energy space environment on the ultraviolet transmittance of optical materials between 1050 A and 3000 A. *Appl Opt* 1966; 5(6): 937-43.

[2] Fröhlich C. Evidence of a long-term trend in total solar irradiance. *Astronomy and Astrophysics* 2009; 501(3).

[3] Gueymard CA, Myers D, Emery K. Proposed reference irradiance spectra for solar energy systems testing. *Solar Energy* 2002; 73(6): 443-467.

[4] Gueymard CA. The sun's total and spectral irradiance for solar energy applications and solar radiation models. *Solar Energy* 2004; 76(4): 423-453.
